# Revisiting the phosphotyrosine binding pocket of Fyn SH2 domain led to the Identification of novel SH2 superbinders

**DOI:** 10.1101/2020.10.30.361790

**Authors:** Shuhao Li, Yang Zou, Dongping Zhao, Yuqing Yin, Jingyi Song, Ningning He, Huadong Liu, Dongmeng Qian, Lei Li, Haiming Huang

## Abstract

Protein engineering through directed evolution is an effective way to obtain proteins with novel functions with the potential applications as tools for diagnosis or therapeutics. Many natural proteins, largely antibodies as well as some non-antibody proteins, have undergone directed evolution in vitro in the test tubes in the laboratories around the world, resulted in the numerous protein variants with novel or enhanced functions. In this study, we constructed a Fyn SH2 variant library by randomizing the 8 variable residues in its phosphotyrosine (pTyr) binding pocket. Selection of this library by a pTyr peptide from MidT antigen led to the identification of SH2 variants with enhanced affinities to the peptide, compared to the wild type SH2, by EC50 assay. Fluorescent polarization (FP) was then applied to quantify the binding affinity of the newly identified SH2 variants. As a result, three SH2 variants, named V3, V13 and V24, have comparable binding affinities with the previously identified SH2 triple-mutant superbinder (refer to Trm). Biolayer Interferometry (BLI) assay was employed to disclose the kinetics of the binding of these SH2 superbinders, in addition to the wild type SH2, to the phosphotyrosine peptide. The results indicated that all the SH2 superbinders have two-orders increase of the dissociation rate when binding the pTyr peptide while there was no significant change in their associate rates. The previously identified SH2 superbinder Trm as well as the V13 and V24 discovered in this study have cross-reactivity with the sulfotyrosine (sTyr) containing peptide while the wild type SH2 does not. Intriguingly, though binding the pTyr peptide with comparable affinity with other SH2 superbinders, the V3 does not bind to the sTyr peptide, implying it binds to the pTyr peptide with a different pattern from the other superbinders. The newly identified superbinders could be utilized as tools for the identification of pTyr-containing proteins from tissues under different physiological or pathophysiological conditions and may have the potential in the therapeutics.

## Introduction

Directed evolution is a process to alter or optimize protein functions [1]. Natural proteins have undergone natural selection to achieve the optimal functions for the physiological process. Nonetheless, only an infinitesimal fraction of the protein sequence space have been explored by natural proteins [2]. It is very attractive to exploit directed evolution to optimize protein functions or acquire novel functions[3–6]. There are two major steps in directed evolution, 1) Mutation of a parental protein, usually on its interaction surface, to obtain a variant library, mostly presented on the surface of a display system(eg. phage, yeast, etc), as the source of the novel proteins. 2) Screen of the library by selective pressure with the purpose of getting protein variants with anticipated functions. For example, a synthetic antibody library is usually constructed by using a natural antibody as a template/scaffold. The CDR(Complementarity Determining Region) randomization of the scaffold antibody results in a synthetic antibody library, in which antibodies for diverse antigens could be selected out[7, 8]. In addition, protein engineers have utilized the non-antibody scaffolds to explore the possibility of obtaining proteins to bind diverse ligands other than the cognate ones, such as Fibronectin type III domain, which is an evolutionary conserved domain of the extracellular protein fibronectin. By diversifying its three loops, i.e. BC, DE and FG loops, protein engineers were able to construct a FN3 variant library. Multiple FN3 variants with high affinity and high specificity to their respective targets have been reported[3–5]. This method was termed monobody technology and was subsequently adopted by biotechnology industry[6]. Similarly, the DARPin and Affibody technologies make use of ankyrins and the Z domain of protein A, respectively, to generated antibody mimetic proteins for the diagnostic and therapeutic applications [9, 10].

In addition to the extracellular proteins mentioned above, intracellular proteins have been harnessed as the scaffolds for proteins with novel functions. For example, the SH3(Src Homology 3)domain of Fyn kinase were randomized at its Src and RT loops and resulted in a Fyn SH3 variant library [11]. The D3 variant, which targets extra-domain B of fibronectin, was selected from this library. Therefore it opened up new biomedical opportunities for the in vivo imaging of solid tumors and for the delivery of toxic agents to the tumor vasculature[11]. Ubiquitin is a 76 amino acids polypeptide relating to protein degradation by the proteasome system, in which the target proteins are ubiquitinatedlabelled ubiquitins by the E1/E2/E3 cascade reaction. The ubiquitin was randomized on its binding surface of its cognate ligands to acquire novel functions, i.e. as inhibitors or activators of the targets in the proteasome system, thereby to be utilized as tools to manipulate the protein degradation process [12]. This allowed the development of potential new therapeutics as well.

Besides obtaining novel functions from parental proteins, directed evolution is also widely used to get protein variants with enhanced functions, usually higher binding affinity. For example, in antibody in vitro affinity maturation, the antigen binding surface, i.e. the CDR region, is usually randomized by site directed mutagenesis to construct a library[13–16]. With affinity selection by its cognate antigen, variant antibodies with more than 10-fold increasing in affinity could be usually obtained [17, 18]. Moreover, intracellular proteins could also be engineered to get high affinity binders to their cognate ligands. For example, SH2 (Src Homology 2)domain is a modular domain that binds selectively to the phosphotyrosine(pTyr)-containing peptides in its cognate binding proteins[19] in the cell signaling pathway with an affinity in the micromolar range [20]. In a previous study, an SH2 superbinder was identified from a Fyn SH2 variant library, in which 15 amino acid residues in the SH2 pTyr binding pocket were randomized[20]. The superbinder, which has an affinity to the phosphotyrosine (pTyr) containing peptides in the single digit nanomolar range, was subsequently applied as a tool to enrich pTyr-containing peptides/proteins, comparing favorably to the conventional anti-pTyr antibodies [21]. In addition, the superbinder not only achieved the function of binding to pTyr peptides with an enhanced binding affinity, but also gained a capacity of binding to sulfotyrosine [21]. Furthermore, the Fyn SH2 superbinder as well as its counterparts (Grb2 SH2&Src SH2) inhibited the EGFR pathway when expressed in vivo, bearing the potential of therapeutic reagents [20][22]. When looking into the selected SH2 variants from the library, we noticed that 7 of the 15 amino acid residues are invariable, showing they are highly conserved during the directed evolution imposed by the selection pressure from the pTyr-containing peptides. In this study, *we* randomized the rest 8 variable residues in the pTyr binding pocket of Fyn SH2 to minimize the theoretical library diversity with the aim to select extra SH2 variants with yet tighter binding affinities to the pTyr peptides. Over one billion of Fyn SH2 variants displayed at the surface of M13 bacteriophages with the designed mutation were biopanned by a pTyr-containing peptide with the sequence of EPQpYEEIPIYL. After four rounds of selection, we identified three more pTyr peptide superbinders, two of which share the sequence features with the previously identified Fyn superbinder Trm. The third variant, named V3, has distinct sequence features. Interestingly, variant V3, though binding to phosphotyrosine with the comparable affinity with the original superbinder, does not have the ability binding to sulfotyrosine in ELISA assay, indicating different mutation pattern in the pTyr binding pocket could confer the variant distinct function.

## Results

### The construction of an SH2 variant library and quality control by next generation sequencing

To construct an SH2 variant library, we randomized the 8 residues(i.e. αA3, BC1, BC2, BC3, βC1, βC3, βD3, βD6 hereafter referred to position 1 to 8, respectively) (Figure 1a and Figure S2) in the Fyn SH2 domain pTyr binding pocket by Kunkel method[23, 24]. Three oligos, which covered the above 8 residues in the template SH2 domain in three regions, were designed for mutation (Figure 1a). In each position, the oligos for Kunkel reaction was doped to bias to the wild type nucleotides(see Material&Methods), resulting in the translated residue as wild type amino acid at approximately 50%, while the other 19 amino acids share the rest 50% (Table S1). The Kunkel reaction was conducted as described [25] and in the Material&Methods section. The resulting Kunkel products were transformed into E. coli SS320, which were pre-infected by helper phage M13KO7 for phage packing, using electroporation. The diversity of the constructed phage display Fyn SH2 variant library was 1.27×10^9^ as determined by clone titration.

**Figure 1.**
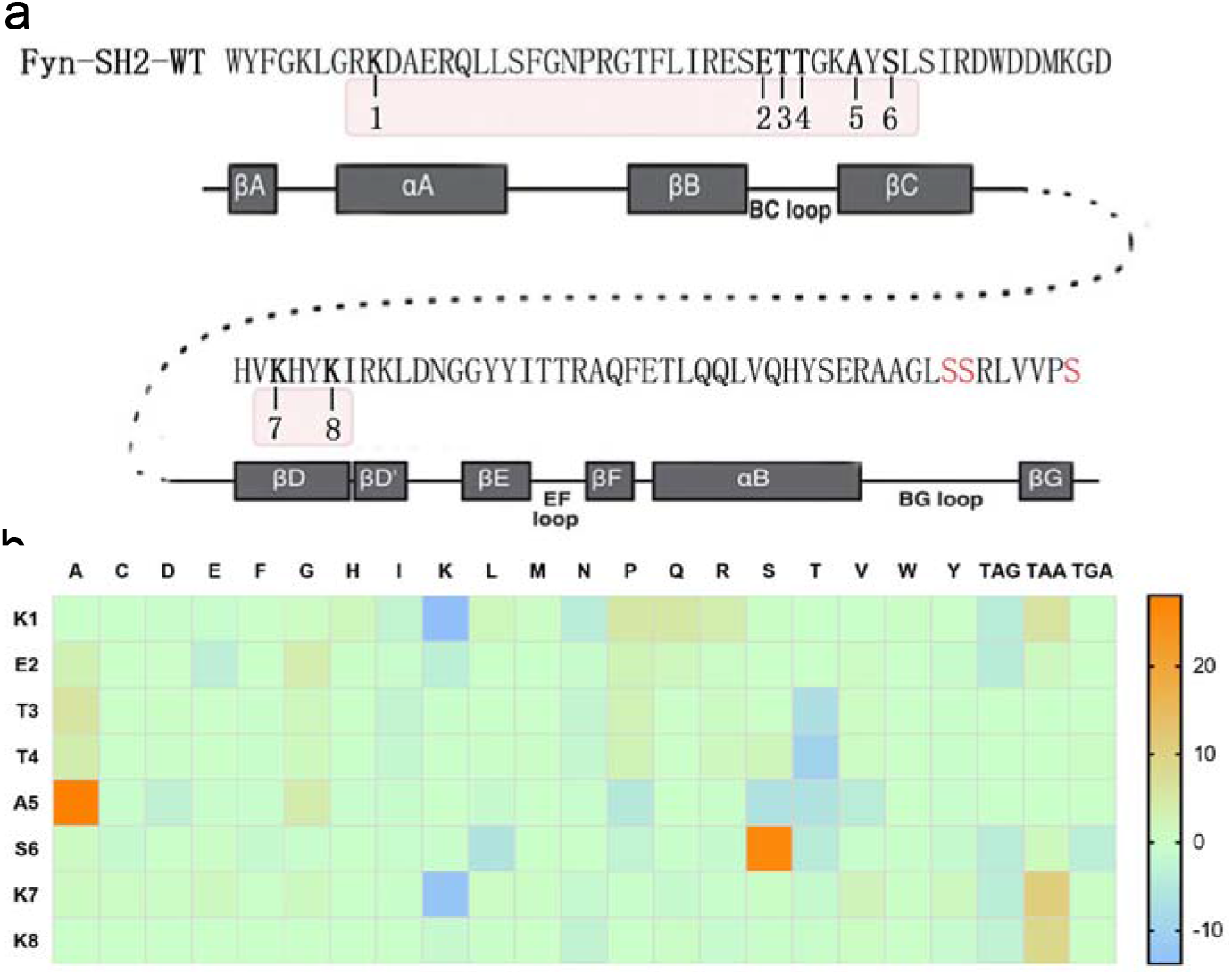
The construction of a Fyn SH2 domain variant library and quality control. a) Amino acid sequence and secondary structure of Fyn SH2 domain. The 8 residues for mutation were numbered. Mutagenesis were introduced by Kunkel method by three primers, in which primer 1 targeting residue 1 (Region 1), primer 2 targeting residues 2,3,4,5 and 6 (Region 2) and primer 3 targeting residue 7 and 8(Region 3) to generate the “stop template”. Then three degenerated primers, ie. primer4(targeting Region 1), prime5(targeting Region 2) and primer6(targeting Region 3), were applied to introduced combinatorial mutations at Region 1, 2 and 3, respectively to generate the library based on the “stop template”. b) The difference between the actual amino acid distribution based on the deep-learning screening and the theoretical amino acid distribution at the 8 positions were shown in the heat map.

To further characterize the library, we used PCR to amplify the coding DNA sequences of the Fyn SH2 variants and subject for deep sequencing. As a results, 5.9 million high quality DNA sequences coding Fyn SH2 variants were retrieved from the deep sequencing results (Table S2). Of these sequences identified, 3.6 million (~60%) have the designed mutations at those 8 positions while the rest have unexpected mutations beyond the 8 positions, likely due to the mutations introduced by the PCR amplification and/or sequencing errors. Among the ~3.6 million sequences with designed mutations, about 55% (2 million) are unique when translated into amino acids. Further analysis of the amino acid distribution of the 5.9 million Fyn SH2 variants showed that the actual mutation of the variant library according with the theoretical design with minor exceptions as show in Figure 1b and Table S1

### Library panning against a pTyr peptide

The library was subject to phage biopanning against a biotinylated peptide EPQpYEEIPIYL derived from protein MidT, which is a cognate ligand of the wild type Fyn SH2[26]. The library phage was pre-incubated with a non-phosphorylated peptide EPQYEEIPIYL, which was immobilized in the wells of a Maxisorp microplate pre-coated with streptavidin. The phage supernatant was then transferred to the wells with the target peptide EPQpYEEIPIYL to enrich the binding phages. Bound phages were eluted and *E.coli XL1-Blue* were infected by the eluted phages for amplification. The amplified phages were applied as the input for the next round of panning. After 4 rounds of panning, the amplified phage pools from each round were applied in an ELISA assay to test their binding to the EPQpYEEIPIYL peptide (see Material & Methods for details). As indicated in Figure 2a, starting from round 3, the phage pools bound specifically to the phosphotyrosine peptide, but not the non-phosphorylated one, indicating the enrichment of the pTyr-specific binding phages.

**Figure 2.**
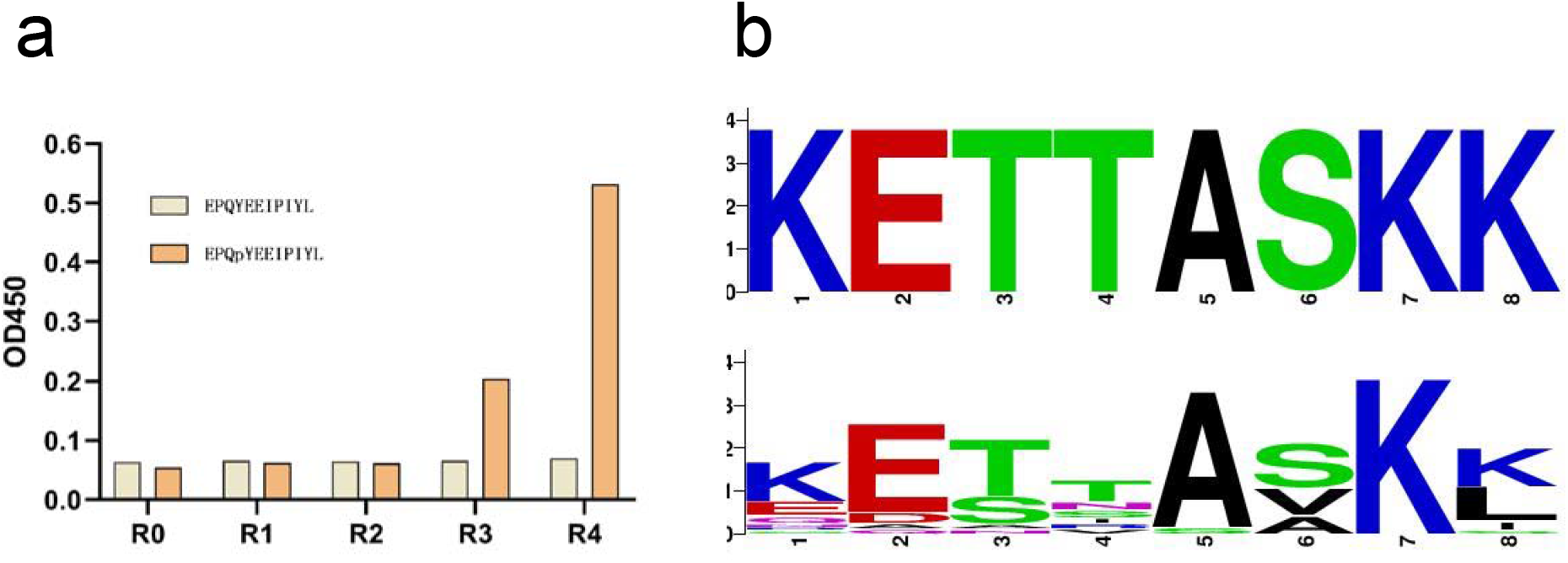
Biopanning of the Fyn SH2 variant library to enrich the pTyr binders. a) Phage ELISA of the selected phage from each biopanning round and the naïve library binding to the phosphotyrosine peptide (EPQpYEEIPIYL) and the non-phosphorylated counterpart, respectively. b) The sequences of ELISA confirmed single SH2 variant clones (n=22) were aligned and the sequences logo of the 8 positions was generated. The sequence of the 8 positions of wild type was also shown at the top for comparison.

To separate single clones that specifically bind to the pTyr peptide, we infected *E.coli XL1-Blue* by phage solution from the enriched rounds (i.e. round 3 and round 4) and plated them on the Carb-positive plate. 192 single bacteria colonies were picked and phage solution were generated by M13KO7 infection. Those 192 phage solution were applied in ELISA to confirm their bindings to pTyr peptide EPQpYEEIPIYL. The clones with OD_450_ ratio (pTyr/non-pTyr) greater than 5 were marked as positive binders and subject to DNA sequencing. It turned out that 22 unique clones, including the wild type Fyn SH2, were identified. We retrieved the 8 residues from each of those 22 clones and generated multi-alignment logo by the online tool WebLogo(https://weblogo.berkeley.edu/). The logo in Figure 2b illustrated that K7 and E2 is very conserved, implying that they play crucial roles to form the structure of the SH2 domain and/or maitain the pTyr peptide binding function. S3 could only be replaced by Threonine and A5 could only be replaced by Glycine. K8 either retains the wild type residue Lysine or adapts to residue Leucine, except one variant has Isoleucine at this position. The mutation scope of S6 is restricted to hydrophobic residues Alanine and Valine. K1 and T4 are the most variable positions as they can be replaced by more than 5 different residues other than the wild type ones.

We then purified all these 22 variants as well as the previously identified superbinder SH2 as His-tag proteins and quantified their binding abilities to EPQpYEEIPIYL peptide by EC50 assay (Table 1 and Figure S1). We classified the clones into three groups based on their EC50 value, i.e. the low affinity (EC50>1000nM) variants, moderate affinity (500nM<EC50<1000nM) variants and high affinity (EC50<500nM) variants. We firstly noticed that the single mutation at K8L (i.e. the variant V1) is enough to increase the binding affinity of the SH2 domain to the pTyr peptide. The variant V10 (T4V/K8L), which has a further mutation at position 4, binds to the pTyr peptide even tighter, with a EC50 value of 175nM. The synergistic effect of the mutations at those two position was also reported previously [20]. The triple mutant with the mutation T4V/S6A/K8L, which has an additional mutation t position 6, was identified as a superbinder previously [20]. In this study, the EC50 value of the triple mutant is 8-fold (21nM) smaller than that of the double mutant variant V10 (T4V/K8L), which verified that the triple mutant is a superbinder to the pTyr peptide and confiremed the synergistic contribution of S6A mutation with T4V/K8L mutation. The contribution of K8L to the binding has been exemplified in the previous study as it is located in the center of a hydrophobic patch formed by the T4V/S6A/K8L residues [20]. We noticed that among those 22 variants, 9 of them have K8L mutation and their binding affinities to the pTyr peptide all increased as the EC50 values ranged from 21nM to 500nM. For those 9 variants, if mutation at positon 4 and/or position 6 added as in the case of the triple mutant, the higher affinities achieved, e.g. V13, V24, etc. However, if the mutations occurred only at the first three positions, either single or double mutation, i.e. V16, V18, V21, V23, those variants we got in this study didn’t show affinity increase at all. As seen in Table 2, there are 6 variants (ie V2, V4, V6, V7, V19 and V22) have T6V mutation at position 6. Although 5 of them (except for V2) have mutation at position 4, which is similar to the triple mutant, the binding affinities of them didn’t increase either. The reason might be that those mutants don’t contain K8L mutation at position 8, which is crucial for the high affinity binding. The fact that T6V and K8L don’t co-exist in one variant implies that T6V mutation and K8L mutation exclude one another during the *in vitro* evolution.

**Table 1.**
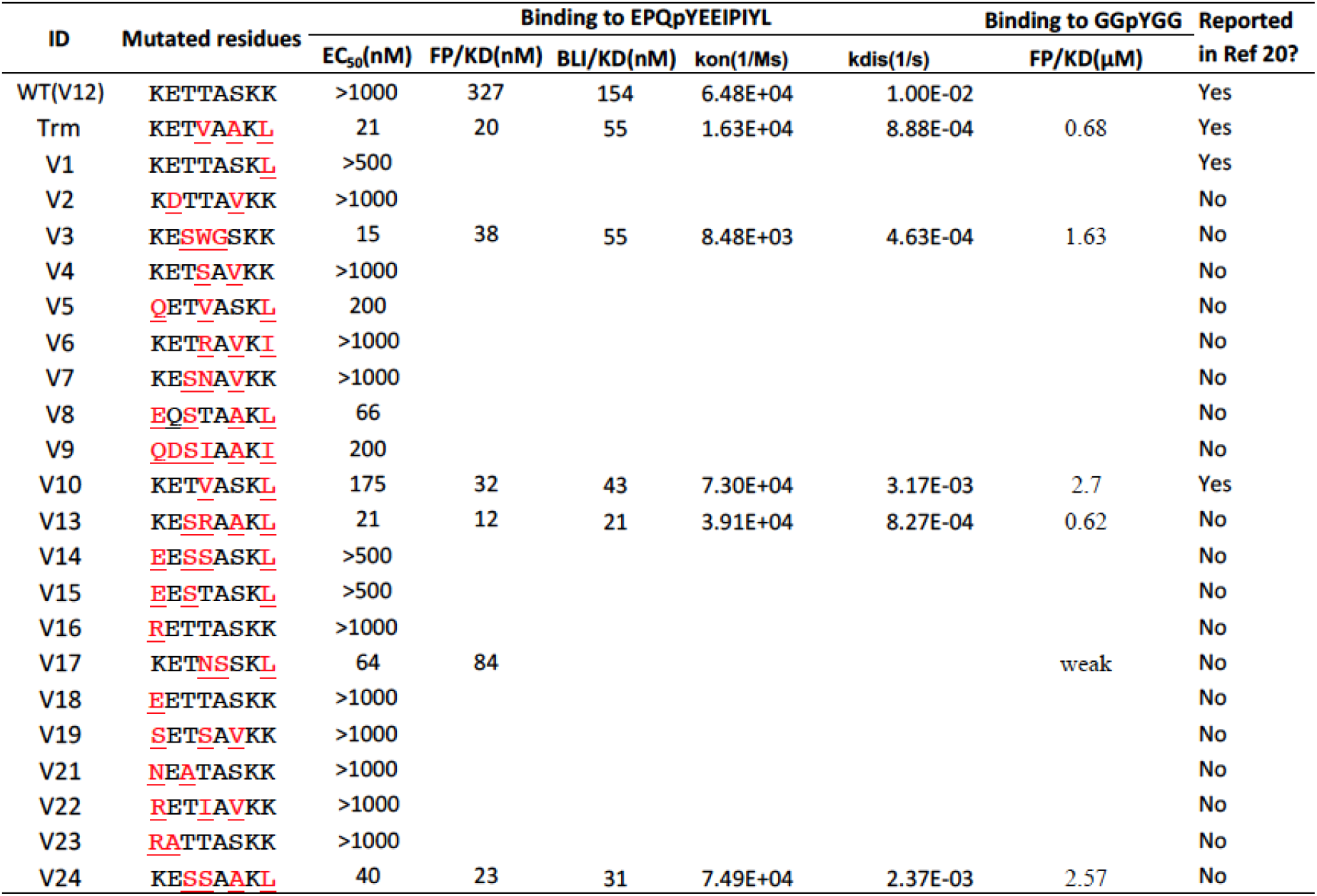
The amino acid residues at those 8 positions and the binding affinities of Fyn SH2 variants to phosphotyrosine peptides measured by different methods.

Variant V3 has mutations at position3 (T3S), position4 (T4W) and position 5(A5G). Initially, the DNA sequencing results showed that there was an ochre stop codon TGA at the position 4, implying an invalid translation/display of the full length Fyn SH2 variant. However, as there were more than 10 clones sharing the same mutations (i.e. T3S/T4[ochre stop codon]/A5G) and were all ELISA positive, suggesting this TGA must be translated into a certain amino acid in this case. We expressed and purified the variant V3 protein and submitted it for mass spectrometry assay identification. The result indicated that the TGA was translated into residue Tryptophan (Figure 3a), so we decoded TGA codon into tryptophan in this case. Furthermore, the EC50 of V3(T3S/T4W/A5G) is 15 nM(Figure 3b), which is very close to the triple mutant (EC50=21nm). Therefore we identified a new mutation pattern, i.e. position 3/4/5 which also resulted in high affinity Fyn SH2 variant to the EPQpYEEIPIYL peptide, in addition to the previously identified pattern positioning in the K8L as well as the K8L related variants as shown above.

**Figure 3.**
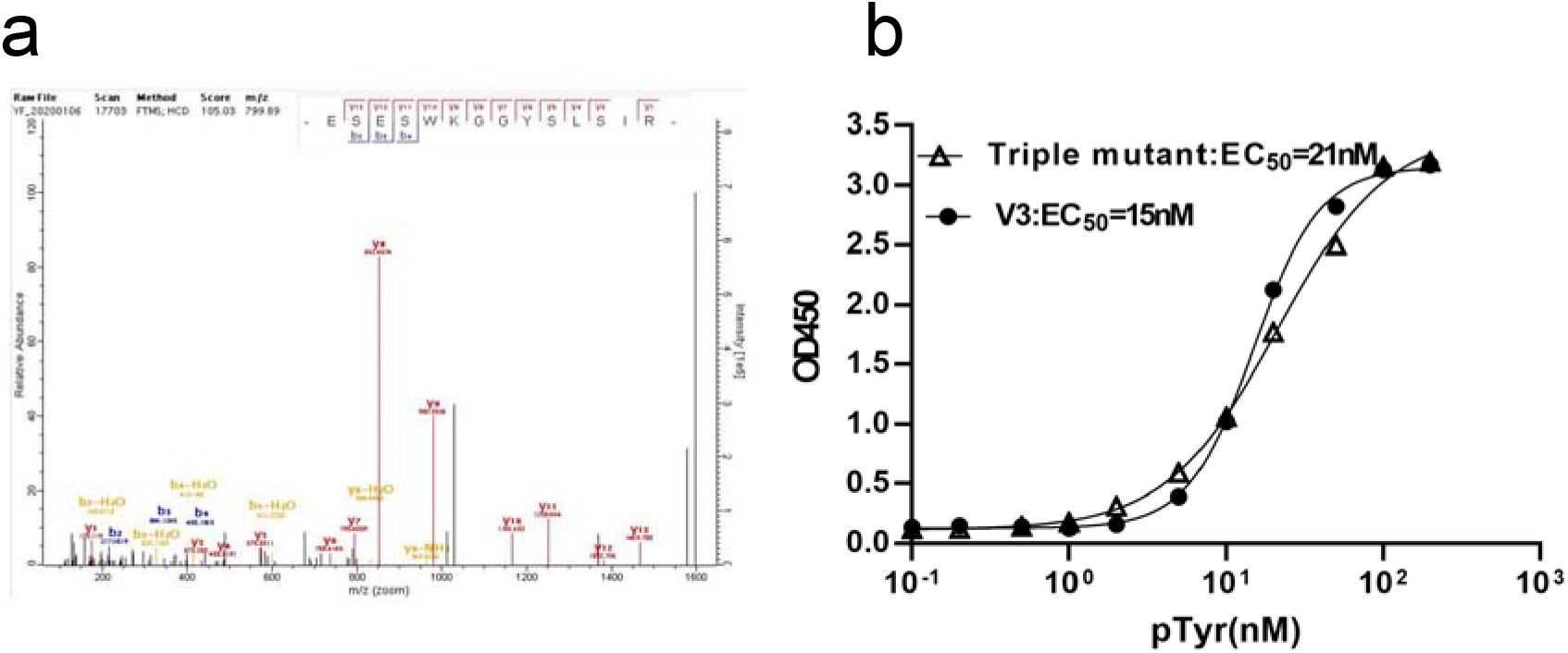
Decoding of the opal stop codon UAG and its role in a functional SH2 variant. a) The V3 variant was expressed and purified. The sample was trypsin digested and submitted for mass spectrometry assay. b) The EC50 assay of the V3 and triple mutant variants binding to the phosphotyrosine peptide (EPQpYEEIPIYL).

### Identification of new superbinders based on binding affinity

To determine which variants were new superbinders in our study, we measured the binding affinities of of those variants that had low EC50 (<200nM) with EPQpYEEIPIYL peptide by FP(Fluorescence polarization). As shown in Table 2 and Figure 4a, the wild type Fyn SH2 binds to the pTyr peptide with a KD of 327nM, which was similar to the result of the previously study[20]. The triple mutant SH2 had a KD of 20nM (Figure 4b), which was also in the affinity range of the mutant binding to the pTyr peptides reported previously [20]. All the other variants from this study had tighter binding affinity to the pTyr peptide than the wild type Fyn SH2 does. The variant V10 (T4V/K8L), also identified in the previous study [20], had a binding affinity of 32nM (Figure 4c), which was slightly weaker than the triple mutant. This is also consistent with the previous data [20]. Variant V17(T4N/A5S/K8L) has an unusual threonine to asparagine mutation and alanine to serine mutation at position 4 and 5, respectively, resulting in a 4-fold affinity increase to the wild type Fyn SH2 domain, whereas 4-fold weaker than the triple mutant (Figure 4d). Variants V13(T3S/T4R/S6A/K8L) and V24(T3S/T4S/S6A/K8L) have the same S6A/K8L mutation as the triple mutant(T4V/S6A/K8L) does, meanwhile completely different type of residues at position 4, i.e. positive charged(T4R) and hydrophilic(T4S) residues, respectively. However, variant V13 has a KD of 12nM (Figure 4e) which is slightly tighter than the triple mutant meanwhile variant V24 has a KD of 23nM(Figure 4f) which is very close to the triple mutant. In addition, variant V3 (T3S/T4W/A5G) has an KD of 38 nM (Figure 4g), which is 8.6-fold tighter than the the wild type Fyn SH2 domain, although is slightly weaker than the triple mutant. So far, we have identified 5 variants (V10, V17, V13, V24 and V3) from this study that have high affinity as their binding affinities to peptide EPQpYEEIPIYL are under 40nM. To verify if they can bind the pTyr moiety alone like the triple mutant, we synthesized an artificial peptide GGpYGG and tested its binding affinities to the above 5 variants by Fluorescence polarization, respectively. As indicated in Table 1 and Figure 5, the triple mutant binds to the GGpYGG peptide with a KD of 0.68μM, which is consistent with the previous result(0.71μM) [20]. Variant V10 (T4V/K8L) had a binding affinity of 2.7μM with peptide GGpYGG *versus* 3.1μM in the previous study [20]. Variants V13 (T3S/T4R/S6A/K8L) and V24 (T3S/T4S/S6A/K8L) had the binding affinities of 0.62 μM and 2.57 μM, respectively, likely because that they share the same S6A/K8L with the triple mutant. Surprisingly, variant V17 (T4N/A5S/K8L) had almost no binding to GGpYGG peptide. In the previous study, variant with the K8L mutation bound to the GGpYGG peptide with the affinity of 13μM[20]. It is likely that the T4N/A5S mutation is deteriorative to the pTyr binding pocket in terms of binding to the pTyr moiety. Interestingly, variant V3(T3S/T4W/A5G), though having the position 4&5 mutation, bound to the GGpYGG peptide with an affinity of 1.63μM. Based on the binding affinity data, we named the variant V13, V24 and V3 the newly identified SH2 superbinders.

**Figure 4.**
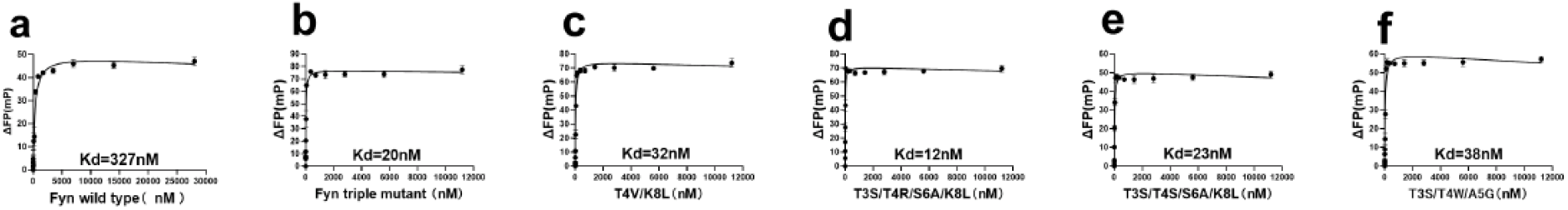
Binding affinities of the SH2 variants to the phosphotyrosine peptide (EPQpYEEIPIYL) measured by FP, including the wild type (a), the triple mutant (b), the variants V10 (c), V17 (d), V13(e), v24 (f) and V13 (g).

**Figure 5.**
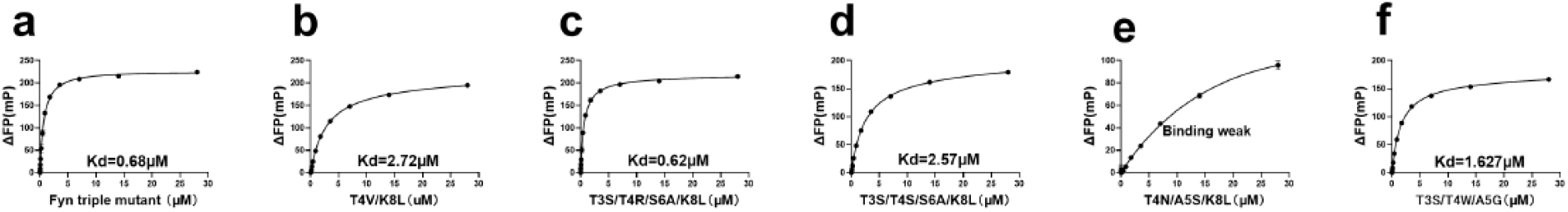
Binding affinities of the SH2 variants to phosphotyrosine moiety (GGpYGG) measured by FP, including the wild type (a), the triple mutant (b), the variants V10 (c), V17 (d), V13(e), v24 (f) and V13 (g).

### Superbiners have slower dissociation rates

To further understand how those superbinders obtained the super binding capability kinetically, we measured their association and dissociation rates binding to the EPQpYEEIPIYL peptide by Biolayer Interferometry (BLI) assay while using the wild type Fyn SH2 as a control. As seen in the Figure 6 and Table 1 the wild type Fyn SH2 has a *k_on_* rate of 6.48E+04 Ms^−1^, which is comparable to most conventional antibodies binding to their antigens. However, the *K_off_* is 1.00E-02 s^−1^, resulting in a measured KD of 154nM, which is relatively strong for the physiologic interactions. Interestingly, the triple mutant has a *k_on_* rate of 1.63E+04 Ms^−1^. Though almost 4-fold slower than the wild type Fyn SH2, it has a two order decrease in *k_off_* rate, which is 8.88E-04 s^−1^, leading to a measured KD of 55nM. The increased *k_off_* rate is consistent with the fact that the T4V/S6A/K8L mutation in the triple mutant created a hydrophobic patch engaging the aromatic ring of the pTyr moiety with hydrophobic interaction[20]. Similarly, both the newly identified variants V13 and V24 in this study have close *k_on_* rate as the wild type domain does, but much slower *k_off_* rate. This is likely due to the S6A/K8L combinatorial mutation that creates the extra hydrophobic interaction with the pTyr aromatic ring, even though they have positive charge residue, arginine, and hydrophilic residue, serine, at position 4, respectively. Conversely, the variant V3 has a 7-fold slower *k_on_* rate than that of the wild type SH2 domain, which is 8.48E+03 Ms^1^. However the *k_off_* rate of V3 (T3S/T4W/A5G) is 4.63E-04 s^−1^, making its binding affinity as high as the triple mutant(T4V/S6A/K8L), i.e. 55nM. The different mutation combination in the protein primary sequence and the *k_on_* rate of V3 imply that it binds to the pTyr peptide in a different mode from the triple mutant(T4V/S6A/K8L).

**Figure 6.**
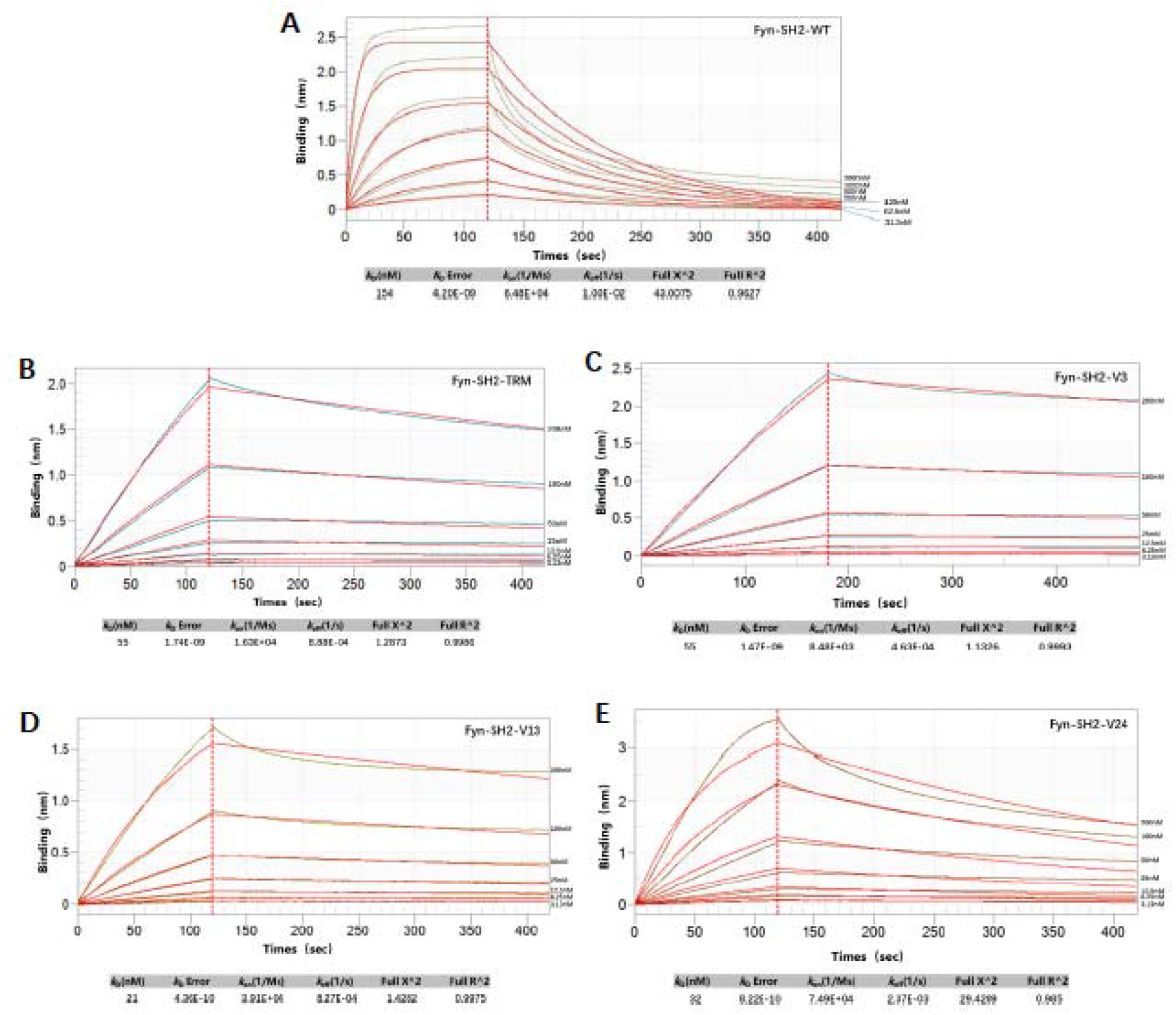
Biolayer Interferometry (BLI) assay to measure the binding affinity of phosphotyrosine peptide(EPQpYEEIPIYL) binding to a)wild type Fyn SH2, b)SH2 triple mutant(TRM), c)variant V3, d)variant V13, e)variant V24.

### Variant V3 has not only high affinity but also high specificity to the pTyr peptide

It was reported that the triple mutant(T4V/S6A/K8L) its variants can bind to sulfotyrosine peptide, in addition to the phosphopeptide [21]. To test if the superbinders discovered in our study could also have this dual function, we prepared the phage solutions that displaying the triple mutant SH2 domain, variant V13 and variant V3, respectively. In a phage ELISA assay, the variant V13, behaving similarly to the triple mutant SH2 domain, bound both sulfotyrosine peptide (EPQsYEEIPIYL) and phosphopeptide (EPQpYEEIPIYL) (Figure 7a). Interestingly, the variant V3 bound exclusively to the phosphopeptide, but not to the sulfotyrosine peptide at all. When expressed as proteins, the variant V13 and V3 showed the similar binding specificities to pTyr and sTyr as measured by protein ELISA (Figure 7b). Based on this finding, we deduced that the T3S/T4W/A5G mutation of variant V3 give rise to not only the interaction with the aromatic ring of the tyrosine, but also discriminating the modification groups (ie. PO_4_^3-^ and SO_4_^2-^) of tyrosine.

**Figure 7.**
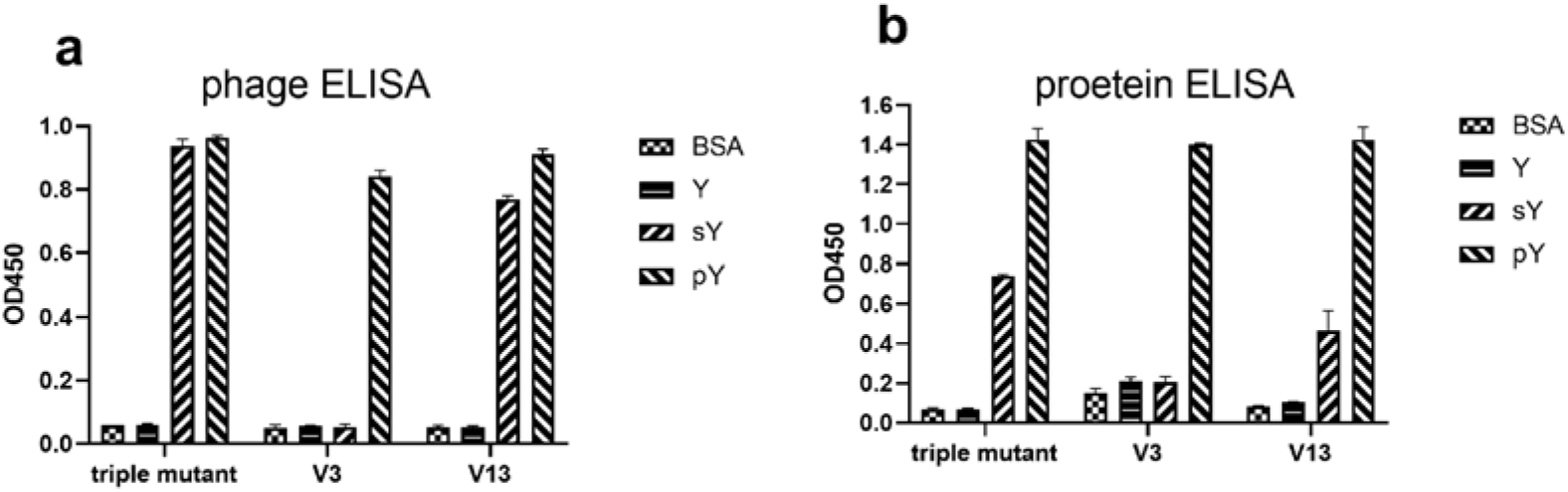
Cross reactivity of wild type Fyn SH2, V3 and V13 binding to phosphotyrosine peptide(EPQpYEEIPIYL) and sulfotyrosine peptide(EPQsYEEIPIYL), respectively, measured by phage ELISA(a) and protein ELISA(b).

## Discussion

In this study, we revisited the pTyr binding pocket of Fyn SH2 domain by randomizing only 8 positions in the pocket. Although the theoretical diversity of the mutation is 2.56X10^10^(20^8^), the actual library size we constructed is 1.29X10^9^ and the expected clones is about 60% as exemplified by DNA deep sequencing. Nonetheless, we were able to got 12 pTyr binders with enhanced binding affinities, likely due to the bias design of this library (See Methods). Soft randomization method, in which the wild type residues have the dominant proportion, is widely used in *in vitro* affinity maturation, especially for antibody engineering [17, 27, 28]. The rationale is that if a residue in the binding pocket contributes less to the binding affinity, the other one(s) will show up during the biopanning by affinity selection, even if this/these residue has/have fewer proportion at this position in the library. So only those residues that contribute more to the binding can compete with the wild type residues and show up in the final selected clones. From the logo generated from the 22 clones selected from this study, we found it is almost identical to the previous study[20], which verified the variability of those 8 positions in the pTyr binding pocket even in a smaller size library. However, among the 22 clones, there are only two clones overlap with those in the previous study(Table 2), implying that various mutation combinations could result in high affinity binders.

Randomizing the key residues which contribute to binding is widely used in construction of library for affinity selecting in order to obtain higher affinity binders, e.g., in vitro affinity maturation in antibody engineering. A few library molecules (from 10 to a few hundred) were usually sent for Sanger sequencing to verify the mutation in the quality control stage. To answer if the actual library we constructed is in line with our design, we submit the whole library for deep sequencing by HiSeq and did the statistical analysis with the purpose of getting a higher resolution of the quality of our library. The NGS data showed that most mutation agreed with the design in spite of a few exceptions. The high quality of the library ensured us getting enough positive clones for the identification of superbinders in the screening stage. Although we got 22 positive clones from the initial biopanning, it didn’t guaranty all of them are higher affinity binders. Despite all this, more than half of them have higher affinity than the wild type Fyn SH2, 3 of which are superbinders as identified by measuring their binding constants. Therefore it is very likely to obtain high affinity binders by selecting a bias library against its cognate ligand. In conclusion, soft randomization is a powerful method to do *in vitro* affinity maturation, in addition to the error prone PCR method.

There are growing evidences to show that stop codon read-through is a common phenomenon in bacteria [29], yeast [30] and human [31]for all three types of canonical stop codons, i.e. UAG, UGA and UAA. Stop codon read-through may be a molecular error [32] or more likely a programmed event in physiological condition[33]. The UGA read-through usually results in the translation of Tryptophan, Cysteine, or Arginine [30]. The *3’* cytosine after UGA facilitates the read-through partially because of the compromised sampling ability of eRF1, which specifically senses cytosine at the +4 position [34, 35]. In our study, we found UGA read-through in the context of -UGAAAA-[Figure S2]. Although not in the ideal context of -UGACU-, it was still translated into tryptophan, but not Cysteine or Arginine, as verified by mass spectrometry. Our finding is also supported by the conclusion from previous study that the -UGAA- tetranucleotide is preferentially read through by tryptophan nc-tRNA [34]. To our best knowledge, our finding is the first report about UGA read-through in a synthetic gene in the phage display system, which may help to rescue those sequences previously discarded in the sequence analysis as they have stop codons. For example, in *in vitro* antibody affinity maturation library design and construction, stop codons are inevitably introduced into the template during library construction either by error-prone PCR or soft randomization used in this study. Our finding could be a heads-up that those sequences with stop codons may be the ture positive clones if enriched during biopanning.

As the UGA is decoded as Trp in variant V3, the T4W mutation may increase the hydrophobicity in the pTyr binding pocket. Furthermore, the T3S and A5G mutations may make more space in the pocket to accommodate the pTyr moiety as those two positions were replaced by two amino acids with smaller side chain. Although the 3D structure of V3 remains resolved, we reasoned that the T3S/T4W/A5G mutation generated increased interaction force between the pTyr moiety and the pTyr pocket while kept the structure of the SH2 variant and the pTyr binding pocket intact. That also makes V3 a different superbinder type from the triple mutant (T4V/S6A/K8L). It is also understandable that that variants V13 (T3S/T4R/S6A/K8L) and V24 (T3S/T4S/S6A/K8L) are superbinders as they share the same mutation pattern with the triple mutant (T4V/S6A/K8L) at position 6 and position 8. Based on the data of these three variants (V13, V24 and the triple mutant), we deduced that the K8L is a crucial mutation to increase the binding affinity as verified in this and the previous studies[20]. We hypothesized that K8L mutation in any Fyn SH2 variant contributes to the binding affinity as verified in this study, i.e. variant V8, V5, V17 from this study, etc., in addition to the V13 and V24.

In summary, we have identified three SH2 superbinders in addition to the first generation triple mutant by engineering the pTyr binding pocket of Fyn SH2 domain. Like the first triple mutant, the newly identified superbinders may be applied in profiling the phosphorylation level of tissue or even bear great promise as antagonists of tyrosine kinase signaling and thereby potential therapeutic agents.

## Material and Methods

### Phage display library construction through site-directed mutagenesis by Kunkel reaction

The wild type human Fyn SH2 domain (Cys-less) in phagemid pFN-OM6 (ref) were used as the template for library construction. The dU-ssDNA of the Fyn SH2 domain as the template in Kunkel reaction was made as described before. Primer 1 (GGAAAATTAGGAAGA*<u>CCATGG</u>*GATGCTGAAAGACAA), Primer 2(CTTATCCGCGAGAGT*<u>CCATGG</u>*AAAGGT*<u>GCTAGC</u>*TATTCACTTTCTATCCGTGAT),and Primer 3(AAAGGAGACCATGTC*<u>TAA</u>*ATTCGCAAACTTGAC) were used in a combinatorial mutation to construct a “Stop template” by the Kunkel method, in which region1 was incorporated with Nco I restriction enzyme recognition site *<u>CCATGG</u>*, region2 contained both Nco I site(*<u>CCATGG</u>*) and Nhe I site(*<u>GCTAGC</u>*) and region3 included the stop codon TAA. After that, Primer 4 (GGAAAACTTGGCCGANNNGATGCTGAGCGACAG), Primer 5 (CTTATCCGCGAGAGTRNNNWWNWWAAAGGTRWWTATMWNCTTTCTATCCGTGAT) and Primer 6(AAAGGAGACCATGTCNNNCATTATNNNATTCGCAAACTTGAC) were applied simultaneously to synthesize the heteroduplex double strand DNA (dsDNA), in which the designed mutations at each position were incorporate into the “Stop template”. In these primers, N was composed of the mixture of 70% of A, 10% of G, T and C, respectively; M represented the mixture of 70% of T and 10% of A, G and C; R related to the mixture of 70% of G, 10% of A, T and C; W contains 70% of A, 10% of T, G and C. The dsDNA was further digested by the restriction enzymes Nco I and Nhe I to remove the unreacted template molecules before the transformation into *E. coli* SS320 (preinfected by M13KO7) by electroporation. The transformation efficiency (library diversity) was calculated by bacterial serial dilution as described [25]. The resulting phage library was precipitated by PEG/NaCl (20% PEG 8000/2.5M NaCl).

### Library quality control by next generation sequencing

Before library preparation, the quality of the DNA samples was assessed on a Bioanalyzer 2100 using a DNA 12000 Chip (Agilent). Sample quantitation was carried out using the Invitrogen’s Picogreen assay. Library preparation was performed according to Illumina’s TruSeq Nano DNA sample preparation protocol. The samples were sheared and uniquely tagged with one of Illumina’s TruSeq LT DNA barcodes to enable library pooling for sequencing. The finished libraries were quantitated using Invitrogen’s Picogreen assay, and the average library size was determined on a Bioanalyzer 2100 using a DNA 7500 chip (Agilent). The Fyn SH2 DNA sequences was amplified by PCR using polymerase chain reaction (PCR) forward primer: 5’-TCCAGGCAGAAGAGTGGTAC-3’ and PCR reverse primer: 5’-AAGTGTTTCAAACTGGGCCC-3’. At last, the library was sequenced on the Illumina HiSeq X™ Ten sequencing system, generating 150 bp paired-end reads.

Paired-end reads were merged using FLASH v1.2.11[36] with default parameters. The merged sequences were quality-filtered by trimmomatic [37] to remove sequences containing more than 3% low-quality bp (Phred score < 3) bases. Sequences were dereplicated, filtered by primer, retaining sequences with identical primers. Then, all the sequences were translated to protein sequences using in-house perl script. Raw sequencing data for this study have been deposited in NCBI Sequence Read Archive database under accession number PRJNA664254.

### Phage biopanning

The phage biopanning was performed as described before [25]. Briefly, 96-well microplate (NUNC, 442404) was coated with 4 pmol streptavidin (Solarbio,S9171) per well in 100μL 1×PBS (137 mM NaCl, 3 mM KCl, 8mM Na_2_HPO_4_ and 1.5mM KH_2_PO_4_, pH=7.2) at 4°C overnight The next morning, the solution in the well was discarded and 200μL/well 0.5% BSA (Bovine albumin, Aladdin, A104912) were added for blocking at room temperature for 1 hour. Biotinylated peptides (biotin-ahx-ahx-EPQYEEIPIYL and biotin-ahx-ahx-EPQpYEEIPIYL), 16pmol/well in 100μL 1× PBS (pH=7.2), were added into two separated wells, labelled as the non-pTyr well and the pTyr well, respectively. After incubation at room temperature for 1 hour, the solution were discarded and 100μL/well phage-displayed SH2 variant library (~1.0×10^11^ phage clones) were added into the non-pTyr well for preclearance for 1 hour. Phage solution were then transferred into the pTyr well for 1 hour. Non-binding phage were washed away by PT buffer (1×PBS+0.05% Tween) for at least 8 times. Bound phages were eluted by 100mM HCl 100μL/well and neutralized by adding 1/8 volume of Tris-HCl (1M, pH=11). Half volume of the neutralized phage solution were then applied to infect 10-fold volume of actively growing *E. coli* XL1-blue (Stratagene) for 30 min at 37°C. Then the M13KO7 helper phage (NEB, N0315S) were added at a final concentration of 1×10^10^ phage/mL for super infection for 45 min. The XL1-blue culture were added into 20-fold volume of 2YT medium (10 g yeast extract, 16 g tryptone, 5 g NaCl in 1L water) supplemented with Carb (carbenicillin, 50 mg/ μL) and Kana (kanamycin, 25 mg/ μL) at 37°C overnight (14-16 hours), 200rpm in a shaker. The overnight culture were centrifuged and the supernatant were precipitated by 1/5 volume of PEG/NaCl (20% PEG 8000/2.5M NaCl). The amplified phage in the pellets were re-suspended with 1mL 1×PBS and applied as the input phage for the next round of panning. From the second round, the immobilized peptide decreased from 16pmol to 12pmol (2^nd^ round), 10pmol (3^rd^ round), 8pmol (4^th^ round) to increase the stringency of the selection.

### Phage ELISA

In a 96-well NUNC microplate, 2 pmol streptavidin were coated per well in 50μL 1×PBS at 4°C overnight. The next morning, the solution in the well was discarded and 100μL/well 0.5% BSA (Bovine albumin, Aladdin, A104912) was added for blocking at room temperature for 1 hour. In the pTyr wells, 8 pmol/well/50μL biotinylated phosphotyrosine peptides (biotin-ahx-ahx-EPQpYEEIPIYL) were added for immobilization for 1 hour at room temperature. Non-phosphrylated peptides were added in the non-pTyr wells as the negative control. The 2×50μL solution was added into the pTyr wells and non-pTyr wells, respectively, for binding for 1 hour. Non-binding phages were washed away 8 times by the PT buffer. 50uL anti-M13/HRP conjugate (Sino Biological, 11973) were added and incubated for 30 min. After wash by the PT buffer, 50uL TMB substrate were added to develop according to the manufacturer’s instruction. 100 μL of 1.0 M H_3_PO_4_ were added to stop the reaction and signals were read spectrophotometrically at 450 nm in a plate reader. The readouts of pTyr and non-Tyr wells were recorded and the ratio of pTyr/ non-Tyr were calculated.

### Protein expression and purification

The cDNA encoding the Fyn SH2 variants in the pFN-OM6 vector were PCR amplified and subcloned into the vector pHH0239 to express 6xHis-tag proteins at the N-terminus. The expression constructs were transformed in to *E. coli* BL21 (DE3). Single colonies were picked and grown in 2YT/Carb medium at 37°C to OD_600_=0.6. IPTG were added to final concentration of 1mM and protein expression was induced at 18 °C overnight. Protein were purified using Ni-NTA agrose (Qiagen, 30210) according to the manufacturer’s manual. The eluted proteins were buffer exchanged into TBS (20 mM Tris-HCl, pH 7.0 and 150 mM NaCl) by Amicon Ultra-4 Centrifugal Filter Units (Millipore). The final concentrations of the proteins were determined by the BCA method.

### EC50 assay

In a 96-well NUNC microplate, 1 pmol SH2 proteins were coated per well in 50μL 1×PBS at 4 °C overnight. 100μL/well 0.5% BSA were added for blocking. A serial biotinylated pTyr peptides with increased concentration (from 0nM, 3.125nM, 6.25nM to 100nM and 200nM) were added in 9 wells coated with SH2 proteins. The wells were washed 4 times by the PT buffer after incubation for 1 hour at room temperature. 50 μL of Streptavidin-HRP conjugate (Sigma, S2438, 1:5000 dilution) were added to each well and incubated for 30 min. After washing 4 times by the PT buffer, 50uL TMB substrates were add to develop color for 2 min. 100 μL of 1.0 M H_3_PO_4_ were added to stop the reaction and signals were read spectrophotometrically at 450 nm in a plate reader. EC50 and standard variation values were calculated using a 3-parameter logistic regression fit using Prism Software (GraphPad).

### Fluorescence polarization binding assay

Peptides were N-terminally labeled with fluorescein. The two 6-aminohexanoic acids (ahx) were used as a linker to couple fluorescein to the peptide. All binding assays were carried out at room temperature in phosphate-buffered saline (PBS) buffer at pH 7.4 and the signals were measured on an VICTOR Multilabel plate reader (Perkin Elmer). Dissociation constants (Kd) were derived from 8 to 12 data points assuming a one-site binding model. Independent measurements (n ≥ 2) produced Kd values within 10% of the reported values. Kd and standard variation values were calculated using Prism Software (GraphPad).

### BLI(Bio-layer interferometry) assay

The BLI experiments were carried out using an Octet RED96 System (ForteBio). The streptavidin biosensors (18-5019) were used to perform the measurement. The biotin-ahx-ahx-EPQpYEEIPIYL peptides were immobilized on the biosensor tip surface. All steps were performed at 30°C with shaking at 1000 rpm in a black 96-well plate (Greiner 655209), with a working volume of 200 μL in each well. The Fyn SH2 variants in the running buffer (1×PBS+0.5% BSA+0.05% Tween) was applied for association for 120 seconds and dissociation for 300 seconds. The response data were normalized using Octet data analysis software version 9.0.0.14 (ForteBio).

## Contribution

LL and HH conceived the project. SL, DZ, YY, JS, NH, HL and DQ conducted the experiments. YZ, LL performed data analysis. HH, LL, SL and DQ wrote the manuscript.

## Acknowledgments

This work was supported in part by funds from the National Natural Science Foundation of China (Grant No. 31770821 and 32071430 to LL); LL is supported by the “Distinguished Expert of Overseas Tai Shan Scholar” program. YZ is supported by the Qingdao Applied Research Project.

---

**Figure S1.**
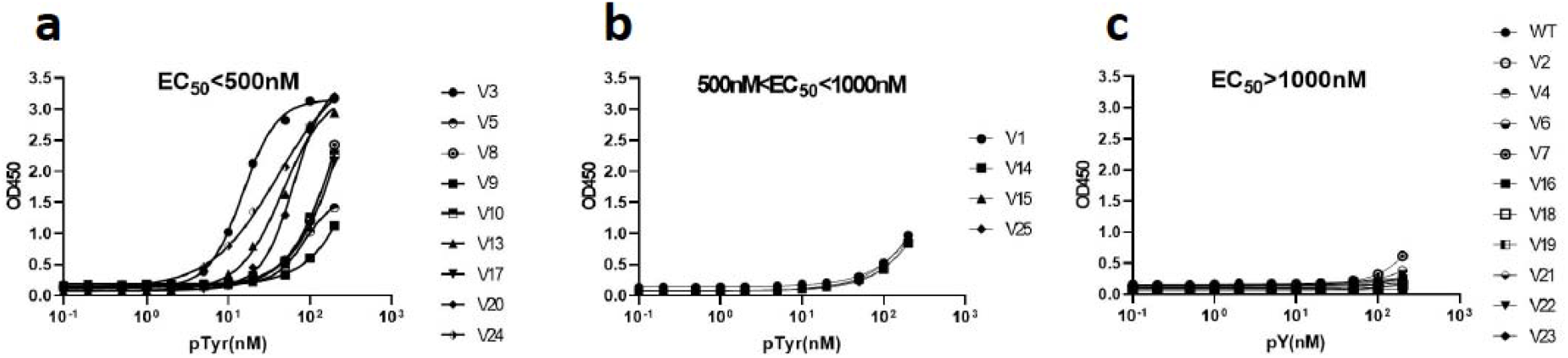
EC50 assay of the 22 SH2 variant proteins selected in this study. a) High affinity binders(EC50<500nM) b) Moderate affinity binders(500nM<EC50<1000nM) c) Low affinity binders(EC50>1000nM).

**Figure S2.**
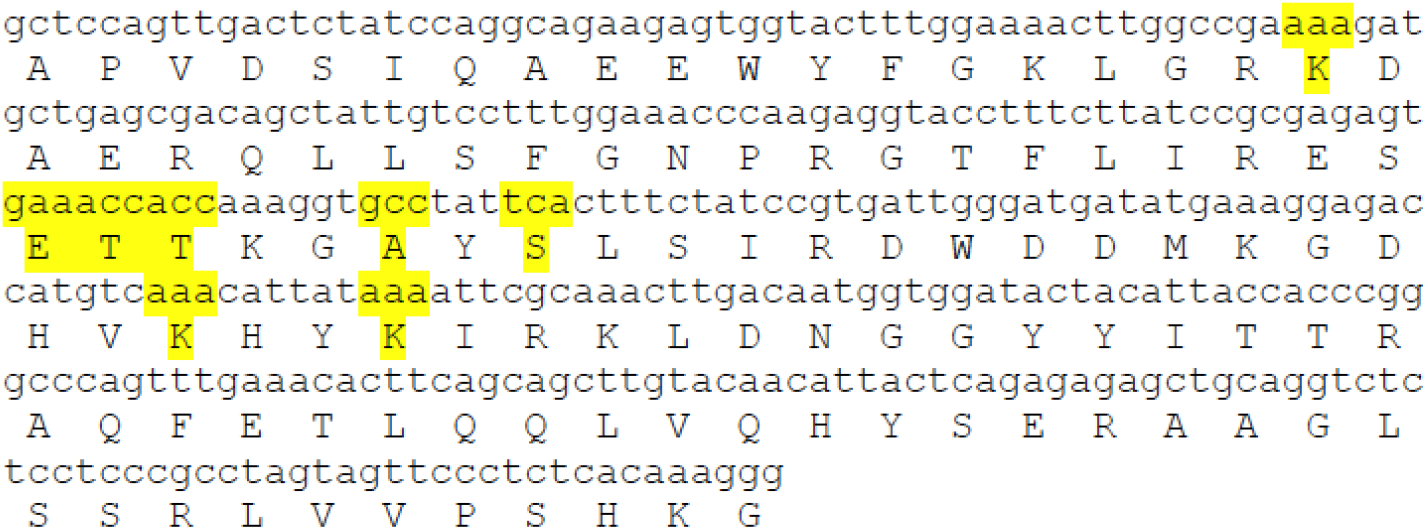
Neucleotide and amino acid sequences of wild type Fyn SH2 domains. The 8 residues for mutated were shaded in yellow.

**Table S1.** Theoretical and actual amino acid distribution at each position.

|  | K1 |  |  | E2 |  |  | T3 |  |  | T4 |  |  | A5 |  |  | S6 |  |  | K7 |  |  | K8 |  |  |
| --- | --- | --- | --- | --- | --- | --- | --- | --- | --- | --- | --- | --- | --- | --- | --- | --- | --- | --- | --- | --- | --- | --- | --- | --- |
|  | Design | Library | Library-Design | Design | Library | Library-Design | Design | Library | Library-Design | Design | Library | Library-Design | Design | Library | Library-Design | Design | Library | Library-Design | Design | Library | Library-Design | Design | Library | Library-Design |
| A | 1.00% | 1.00% | 0.00% | 7.00% | 10.00% | -3.00% | 7.00% | 12.91% | -5.91% | 7.00% | 11.00% | -4.00% | 49.00% | 77.00% | -28.00% | 7.00% | 8.00% | -1.00% | 1.00% | 2.00% | -1.00% | 1.00% | 1.00% | 0.00% |
| C | 0.20% | 0.00% | 0.20% | 0.20% | 0.00% | 0.20% | 0.80% | 1.00% | 0.20% | 0.80% | 1.00% | -0.20% | 0.80% | 0.00% | 0.80% | 1.40% | 0.00% | 1.40% | 0.20% | 1.00% | -0.80% | 0.20% | 0.00% | 0.20% |
| D | 1.40% | 1.00% | 0.40% | 9.80% | 9.89% | -0.09% | 0.80% | 1.15% | 0.35% | 0.80% | 1.06% | -0.26% | 5.60% | 2.72% | 2.88% | 0.20% | 0.10% | 0.10% | 1.40% | 1.99% | -0.59% | 1.40% | 1.25% | 0.15% |
| E | 5.60% | 4.75% | 0.85% | 39.20% | 35.69% | 3.51% | 0.20% | 0.30% | 0.10% | 0.20% | 0.32% | -0.12% | 1.40% | 0.65% | 0.75% | 0.80% | 1.36% | -0.56% | 5.60% | 6.94% | -1.34% | 5.60% | 5.57% | 0.03% |
| F | 0.20% | 0.23% | -0.03% | 0.20% | 0.06% | 0.14% | 0.80% | 0.43% | -0.37% | 0.80% | 0.52% | 0.28% | 0.80% | 0.01% | 0.79% | 1.40% | 0.09% | 1.31% | 0.20% | 0.25% | -0.05% | 0.20% | 0.00% | 0.20% |
| G | 1.00% | 1.79% | -0.79% | 7.00% | 10.83% | -3.83% | 1.00% | 2.73% | -1.73% | 1.00% | 2.53% | -1.53% | 7.00% | 11.29% | -4.29% | 1.00% | 0.45% | 0.55% | 1.00% | 2.37% | -1.37% | 1.00% | 1.16% | -0.16% |
| H | 1.40% | 3.06% | -1.66% | 1.40% | 1.73% | -0.33% | 0.80% | 0.68% | -0.12% | 0.80% | 0.72% | 0.08% | 0.80% | 0.10% | 0.70% | 0.20% | 0.07% | 0.13% | 1.40% | 1.30% | 0.10% | 1.40% | 1.19% | 0.21% |
| I | 6.30% | 4.13% | 2.17% | 0.90% | 0.25% | 0.65% | 6.30% | 4.00% | -2.30% | 6.30% | 4.50% | 1.80% | 0.90% | 0.03% | 0.87% | 0.90% | 0.10% | 0.80% | 6.30% | 5.91% | 0.39% | 6.30% | 6.12% | 0.18% |
| K | 39.20% | 25.45% | 13.75% | 5.60% | 2.19% | 3.41% | 1.40% | 1.02% | -0.38% | 1.40% | 1.31% | 0.09% | 0.20% | 0.02% | 0.18% | 0.80% | 0.33% | 0.47% | 39.20% | 26.69% | 12.51% | 39.20% | 38.10% | 1.10% |
| L | 1.80% | 3.41% | -1.61% | 1.80% | 1.64% | 0.16% | 1.20% | 1.13% | -0.07% | 1.20% | 1.43% | -0.23% | 1.20% | 0.10% | 1.10% | 6.60% | 0.51% | 6.09% | 1.80% | 2.60% | -0.80% | 1.80% | 1.45% | 0.35% |
| M | 0.70% | 1.18% | -0.48% | 0.10% | 0.08% | 0.02% | 0.70% | 0.82% | 0.12% | 0.70% | 0.95% | -0.25% | 0.10% | 0.01% | 0.09% | 0.10% | 0.03% | 0.07% | 0.70% | 1.13% | -0.43% | 0.70% | 0.78% | -0.08% |
| N | 9.80% | 5.81% | 3.99% | 1.40% | 0.54% | 0.86% | 5.60% | 3.46% | -2.14% | 5.60% | 3.94% | 1.66% | 0.80% | 0.06% | 0.74% | 0.20% | 0.04% | 0.16% | 9.80% | 8.16% | 1.64% | 9.80% | 7.20% | 2.60% |
| P | 1.00% | 5.89% | -4.89% | 1.00% | 3.51% | -2.51% | 7.00% | 9.70% | -2.70% | 7.00% | 9.62% | -2.62% | 7.00% | 1.94% | 5.06% | 7.00% | 4.47% | 2.53% | 1.00% | 0.68% | 0.32% | 1.00% | 0.86% | 0.14% |
| Q | 5.60% | 10.83% | -5.23% | 5.60% | 7.22% | -1.62% | 0.20% | 0.21% | 0.01% | 0.20% | 0.25% | -0.05% | 0.20% | 0.13% | 0.07% | 0.80% | 0.39% | 0.41% | 5.60% | 4.17% | 1.43% | 5.60% | 6.25% | -0.65% |
| R | 6.60% | 10.42% | -3.82% | 1.80% | 2.31% | -0.51% | 2.40% | 3.31% | 0.91% | 2.40% | 3.66% | -1.26% | 1.20% | 0.85% | 0.35% | 1.80% | 0.58% | 1.22% | 6.60% | 6.23% | 0.37% | 6.60% | 5.57% | 1.03% |
| S | 2.40% | 2.79% | -0.39% | 1.20% | 1.24% | -0.04% | 12.60% | 12.72% | -0.12% | 12.60% | 14.66% | -2.06% | 7.80% | 1.31% | 6.49% | 49.20% | 75.43% | -26.23% | 2.40% | 2.80% | -0.40% | 2.40% | 2.14% | 0.26% |
| T | 7.00% | 6.81% | 0.19% | 1.00% | 0.90% | 0.10% | 49.00% | 41.80% | -7.20% | 49.00% | 39.25% | 9.75% | 7.00% | 0.71% | 6.29% | 7.00% | 2.73% | 4.27% | 7.00% | 5.66% | 1.34% | 7.00% | 6.51% | 0.49% |
| V | 1.00% | 0.93% | 0.07% | 7.00% | 7.81% | -0.81% | 1.00% | 1.87% | 0.87% | 1.00% | 1.84% | -0.84% | 7.00% | 2.98% | 4.02% | 1.00% | 0.49% | 0.51% | 1.00% | 3.27% | -2.27% | 1.00% | 1.80% | -0.80% |
| W | 0.10% | 0.61% | -0.51% | 0.10% | 0.28% | -0.18% | 0.10% | 0.30% | 0.20% | 0.10% | 0.28% | -0.18% | 0.10% | 0.06% | 0.04% | 0.70% | 0.29% | 0.41% | 0.10% | 0.39% | -0.29% | 0.10% | 0.17% | -0.07% |
| Y | 1.40% | 1.51% | -0.11% | 1.40% | 0.54% | 0.86% | 0.80% | 0.46% | -0.34% | 0.80% | 0.53% | 0.27% | 0.80% | 0.03% | 0.77% | 1.40% | 0.18% | 1.22% | 1.40% | 2.75% | -1.35% | 1.40% | 0.38% | 1.02% |
| TAG | 4.90% | 1.02% | 3.88% | 4.90% | 0.64% | 4.26% | 0.10% | 0.00% | -0.10% | 0.10% | 0.01% | 0.09% | 0.10% | 0.00% | 0.10% | 4.90% | 0.87% | 4.03% | 4.90% | 1.12% | 3.78% | 4.90% | 2.24% | 2.66% |
| TAA | 0.70% | 6.65% | -5.95% | 0.70% | 1.82% | -1.12% | 0.10% | 0.00% | -0.10% | 0.10% | 0.01% | 0.09% | 0.10% | 0.00% | 0.10% | 0.70% | 2.23% | -1.53% | 0.70% | 11.50% | -10.80% | 0.70% | 9.27% | -8.57% |
| TGA | 0.70% | 0.73% | -0.03% | 0.70% | 0.83% | -0.13% | 0.10% | 0.00% | -0.10% | 0.10% | 0.61% | -0.51% | 0.10% | 0.00% | 0.10% | 4.90% | 1.26% | 3.64% | 0.70% | 1.09% | -0.39% | 0.70% | 0.99% | -0.29% |

**TableS2.**
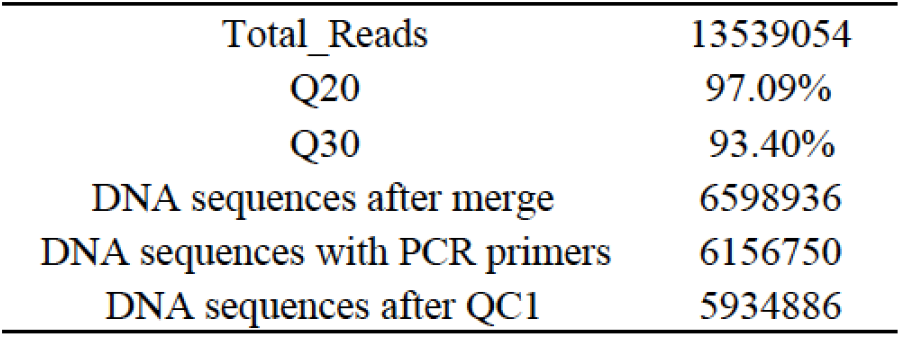
Quality control of the Fyn SH2 variant library by deep sequencing.

|  |  |
| --- | --- |
| Total_Reads | 13539054 |
| Q20 | 97.09% |
| Q30 | 93.40% |
| DNA sequences after merge | 6598936 |
| DNA sequences with PCR primers | 6156750 |
| DNA sequences after QC1 | 5934886 |

